## Supplementary methods for "Polysaccharides induce deep-sea *Lentisphaerae* strains to release chronic bacteriophages"

[*] Corresponding author

Chaomin Sun

### Supplementary Methods

**Transcriptional profiling of *Lentisphaerae* strains WC36 and zth2 cultured in different conditions**

**(1) Library preparation for strand-specific transcriptome sequencing.** A total amount of 3 μg RNA per sample was used as input material for the RNA sample preparations. Sequencing libraries were generated using NEBNext® Ultra™ Directional RNA Library Prep Kit for Illumina® (NEB, USA) according to manufacturer's recommendations and index codes were added to attribute sequences to each sample. The rRNA is removed using a specialized kit that leaves the mRNA. Fragmentation was carried out using divalent cations under elevated temperature in NEBNext First Strand Synthesis Reaction Buffer (5×). First strand cDNA was synthesized using random hexamer primer and M-MuLV Reverse Transcriptase (RNaseH$^-$). Second strand cDNA synthesis was subsequently performed using DNA Polymerase I and RNase H. In the reaction buffer, dNTPs with dTTP were replaced by dUTP. Remaining overhangs were converted into blunt ends via exonuclease/polymerase activities. After adenylation of 3' ends of DNA fragments, NEBNext Adaptor with hairpin loop structure was ligated to prepare for hybridization. In order to select cDNA fragments of preferentially 150~200 bp in length, the library fragments were purified with AMPure XP system (Beckman Coulter, Beverly, USA). Then 3 μL USER Enzyme (NEB，USA) was used with size-selected, adaptor-ligated cDNA at 37 ℃ for 15 minutes followed by 5 minutes at 95 ℃ before PCR. Then PCR was performed with Phusion High-Fidelity DNA polymerase, Universal PCR primers and Index (X) Primer. At last, products were purified (AMPure XP system) and library quality was assessed on the Agilent Bioanalyzer 2100 system.

**(2) Clustering and sequencing.** The clustering of the index-coded samples was performed on a cBot Cluster Generation System using TruSeq PE Cluster Kit v3-cBot-HS (Illumia) according to the manufacturer's instructions. After cluster generation, the library preparations were sequenced on an Illumina Hiseq platform and paired-end reads were generated.

**(3) Data analysis.** Raw data (raw reads) of fastq format were firstly processed through in-house perl scripts. In this step, clean data (clean reads) were obtained by removing reads containing adapter, reads containing ploy-N and low quality reads from raw data. At the same time, Q20, Q30 and GC content the clean data were calculated. All the downstream analyses were based on the clean data with high quality. Reference genome and gene model annotation files were downloaded from genome website directly. Both building index of reference genome and aligning clean reads to reference genome were used Bowtie2-2.2.3 (Langmead and Salzberg, 2012). HTSeq v0.6.1 was used to count the reads numbers mapped to each gene. And then FPKM of each gene was calculated based on the length of the gene and reads count mapped to this gene. FPKM, expected number of Fragments Per Kilobase of transcript sequence per Millions base pairs sequenced, considers the effect of sequencing depth and gene length for the reads count at the same time, and is currently the most commonly used method for estimating gene expression levels (Anders and Huber, 2010).

**(4) Differential expression analysis.** Differential expression analysis of two conditions/groups (two biological replicates per condition) was performed using the DESeq R package (1.20.0) (Wang et al., 2010). DESeq provide statistical routines for determining differential expression in digital gene expression data using a model based on the negative binomial distribution. The resulting *P*-values were adjusted using the Benjamini and Hochberg's approach for controlling the false discovery rate. Genes with an adjusted $P < 0.05$ found by DESeq were assigned as differentially expressed. (For DEGSeq without biological replicates) Prior to differential gene expression analysis, for each sequenced library, the read counts were adjusted by edgeR program package through one scaling normalized factor. Corrected *P*-value of

0.005 and $\log_2$ (Fold change) of 1 were set as the threshold for significantly differential expression.

**(5) GO and KEGG enrichment analysis of differentially expressed genes.** Gene Ontology (GO) enrichment analysis of differentially expressed genes was implemented by the GOseq R package, in which gene length bias was corrected (Young et al., 2010). GO terms with corrected *P* value less than 0.05 were considered significantly enriched by differential expressed genes. KEGG is a database resource for understanding high-level functions and utilities of the biological system, such as the cell, the organism and the ecosystem, from molecular-level information, especially large-scale molecular datasets generated by genome sequencing and other high-throughput experimental technologies (http://www.genome.jp/kegg/) (Kanehisa et al., 2008). We used KOBAS software to test the statistical enrichment of differential expression genes in KEGG pathways.

**Quantitative real-time PCR assay.** For qRT-PCR, cells of strain WC36 were cultured in basal medium supplemented with or without 10 g/L laminarin for 5 days and 10 days, and cells of strain zth2 were cultured in basal medium supplemented without or with 3 g/L laminarin for 4 days. Total RNAs from each sample were extracted using the Trizol reagent (Solarbio, China) and the RNA concentration was measured using Qubit® RNA Assay Kit in Qubit® 2.0 Flurometer (Life Technologies, CA, USA). Then RNAs from corresponding sample were reverse transcribed into cDNA and the transcriptional levels of different genes were determined by qRT-PCR using SybrGreen Premix Low rox (MDbio, China) and the QuantStudioTM 6 Flex (Thermo Fisher Scientific, USA). The PCR condition was set as following: initial denaturation at 95 °C for 3 minutes, followed by 40 cycles of denaturation at 95 °C for 10 s, annealing at 60 °C for 30 s, and extension at 72 °C for 30 s. 16S rRNA was used as an internal reference and the gene expression was calculated using the $2^{-\Delta\Delta Ct}$ method, with each transcript signal normalized to that of 16S rRNA. Transcript signals for each treatment were compared to those of control group. Specific primers for genes encoding phage-associated proteins and 16S rRNA were designed using Primer 5.0 as shown in Supplementary file 6. All qRT-PCR runs were performed in three biological and three technical replicates.

**A detailed procedure for genome sequencing analysis of bacteriophages**

**(1) Library construction and Illumina HiSeq sequencing.** Briefly, for Illumina pair-end sequencing of each phage, 1.0 μg genomic DNA was used for the sequencing library construction. Paired-end libraries with insert sizes of ~ 400 bp were prepared following the standard procedure. The purified genomic DNA was sheared into smaller fragments with a desired size by Covaris, and blunt ends were generated using the T4 DNA polymerase. And the desired fragments were purified through gel-electrophoresis, then enriched and amplified by PCR. The index tag was introduced into the adapter at the PCR stage and we performed a library quality test. Finally, the qualified Illumina pair-end library was used for Illumina NovaSeq 6000 sequencing (150 bp*2, Shanghai BIOZERON Co., Ltd).

**(2) Genome assembly.** The raw paired end reads were trimmed to remove the Illumina adaptors and to retain high-quality reads (score of >30 and length of >36 bases), as recommended by the Trimmomatic (version 0.36, http://www.usadellab.org/cms/uploads/supplementary/Trimmomatic) (Bolger et al., 2014) with parameters (SLIDINGWINDOW: 4:15, MINLEN: 75). Then clean data were obtained and used for further analysis. We have used the ABySS software (http://www.bcgsc.ca/platform/bioinfo/software/abyss) to perform genome assembly with multiple-Kmer parameters. VIBRANT v1.2.1 (Kieft et al., 2020), DRAM-v (Shaffer et al., 2020), VirSorter v1.0.5 (with categories 1 (“pretty sure”) and 2 (“quite sure”)) (Roux et al., 2015) and VirFinder v1.1 (with statistically significant viral prediction: score > 0.9 and $P$-value < 0.05) (Ren et al., 2017) with default parameters were used to identify viral genomes from these assembly sequences by searching against the both cultured and non-cultured viral NCBI-RefSeq database (http://blast.ncbi.nlm.nih.gov/) and IMG/VR database (Camargo et al., 2023). The

GapCloser software (https://sourceforge.net/projects/soapdenovo2/files/GapCloser/) was subsequently applied to fill up the remaining local inner gaps and correct the single base polymorphism for the final assembly results. The completeness of viral genomes was estimated using the CheckV v0.6.0 pipeline (Nayfach et al., 2021).

**(3) Genome Annotation.** For bacteriophages, these obtained genome sequences were subsequently annotated by searching these predicted genes against non-redundant (NR in NCBI, 20180814), SwissProt (release-2021_03, http://uniprot.org) (Consortium, 2021), KEGG (Release 94.0, http://www.genome.jp/kegg/) (Kanehisa et al., 2021), COG (update-2020_03, http://www.ncbi.nlm.nih.gov/COG) (Galperin et al., 2021) and CAZy (update-2021_09, http://www.cazy.org/) (Drula et al., 2022) databases. And the CAZymes were inferred from searches of the NCBI nonredundant (nr) protein database with BLASTP (blast.ncbi.nlm.nih.gov), searches of UniProtKB with HMMer (hmmer.org) and searches of CAZy database with dbCAN tool (Yin et al., 2012; Zhang et al., 2018), using an E-value cutoff of $1\times10^{-6}$ for all three. Finally, all the results with the lowest E-value were marked as “hypothetical protein”.

**Operational taxonomic units (OTUs) analysis.** Briefly, total DNAs from different samples were extracted respectively by the CTAB/SDS method (Murray and Thompson, 1980) and diluted to 1 ng/μL with sterile water, which were used for PCR templates. 16S rRNA genes of distinct regions (16S V3/V4) were amplified using specific primers (341F: 5’-CCTAYGGGRBGCASCAG-3’ and 806R: 5’-GGACTACNNGGGTATCTAAT-3’). The PCR products were purified with a Qiagen Gel Extraction Kit (Qiagen, Germany) following the manufacturer's instructions for libraries construction. Sequencing libraries were generated using TruSeq® DNA PCR-Free Sample Preparation Kit (Illumina, USA) according to the manufacturer's instructions. The library quality was assessed on the Qubit@ 2.0 Fluorometer (Thermo Scientific, USA) and Agilent Bioanalyzer 2100 system. Then, the library was sequenced on an Illumina NovaSeq platform and 250 bp paired-end reads were generated. Paired-end reads were merged using FLASH (V1.2.7, http://ccb.jhu.edu/software/FLASH/) (Magoc and Salzberg, 2011), which was designed to merge paired-end reads when at least some of the reads overlap with those generated from the opposite end of the same DNA fragments, and the splicing sequences were called raw tags. Quality filtering on the raw tags was performed under specific filtering conditions to obtain the high-quality clean tags (Bokulich et al., 2013) according to the QIIME (V1.9.1, http://qiime.org/scripts/split_libraries_fastq.html) quality controlled process. The tags were compared with the reference database (Silva database, https://www.arb-silva.de/) using UCHIME algorithm (UCHIME Algorithm, http://www.drive5.com/usearch/manual/uchime_algo.html) (Edgar et al., 2011) to detect chimera sequences, and then the chimera sequences were removed (Haas et al., 2011). Lastly, Uparse software (Uparse v7.0.1001, http://drive5.com/uparse/) (Edgar, 2013) was used to analyze these sequences. Sequences with ≥97% similarity were assigned to the same OTUs. The representative sequence for each OTU was screened for further annotation. For each representative sequence, the Silva Database (http://www.arb-silva.de/) (Quast et al., 2013) was used to annotate taxonomic information based on Mothur algorithm.

### Supplementary Figures legends

**Figure 2—figure supplement 1. TEM observation of potential phage-like particles extracted from the rich medium supplemented with 10 g/L laminarin alone (A) or 10 g/L starch alone (B).**

**Figure 2—figure supplement 2. TEM observation of the rich medium supplemented with 10 g/L laminarin alone (A) or 10 g/L starch alone (B).**

**Figure 2—figure supplement 3. The morphology of *Lentisphaerae* strain WC36 and its released filamentous bacteriophages.** (A) TEM observation of the morphology of released filamentous bacteriophages and strain WC36 cultured in rich medium supplemented without (panels I-III) or with (panels IV-VI) 10 g/L laminarin for 5, 10, and 30 days, respectively. “Lam” indicate the potential laminarin adhered to bacterial cells. The filamentous phages released from bacterial cells were pointed using red arrows. (B) TEM observation of an ultrathin section of strain WC36 cultured in rich medium supplemented without (panel I) or with (panels II-III) 10 g/L laminarin. White arrows indicate the extrusions or buddings around bacterial cells.

**Figure 3—figure supplement 1. TEM observation of filamentous phages extracted from the supernatant of strain WC36 cell suspension cultured in rich medium supplemented with or without 5 g/L or 10 g/L laminarin (for 5, 10, and 30 days).** Panels I-III show the absence of phages in the supernatant of cell strain WC36 cultivated in rich medium for 5, 10, and 30 days, respectively. Panels IV-VI show the morphology of filamentous phages present in the supernatant of cell strain WC36 cultivated in rich medium supplemented with 5 g/L laminarin for 5, 10, and 30 days, respectively. Panels VII-IX show the morphology of filamentous phages present in the supernatant of cell strain WC36 cultivated in rich medium supplemented with 10 g/L laminarin for 5, 10, and 30 days, respectively. Scale bars, 1 μm.

**Figure 4—figure supplement 1. Transcriptome profiles and RT-qPCR analysis of the genes encoding secretion system related proteins.** (A) Transcriptomics-based heat map showing all up-regulated genes encoding secretion system-associated proteins in *Lentisphaerae* strain WC36. "Rich" indicates strain WC36 cultivated in rich medium; "Lam" indicates strain WC36 cultivated in rich medium supplemented with 10 g/L laminarin. (B) RT-qPCR detection of the expression of genes shown in panel A. (C) Transcriptomics-based heat map showing all up-regulated genes encoding secretion system-associated proteins in strain zth2. "Rich" indicates strain zth2 cultivated in rich medium; "Lam" indicates strain zth2 cultivated in rich medium supplemented with 3 g/L laminarin. (D) RT-qPCR detection of the expression of genes shown in panel C. Three replicates were performed for each condition. The locus tag and its corresponding encoding product were shown with the heat map. The heat map is generated by Heml 1.0.3.3. The numbers in panels A and C represent multiple differences in gene expression (by taking $\log_2$ values).

**Figure 5—figure supplement 1. Comparative genome analysis between the Phage-WC36-1 and Phage-zth2-1.**

**Figure 5—figure supplement 2. Phylogenetic analysis of Phage-WC36-1 and related phages.** (A) Maximum likelihood phylogenetic tree of single-stranded DNA-binding protein sequences from phage-WC36-1 and some related phages. *Chlamydia pneumoniae* J138 was used as the outgroup. (B) Maximum likelihood phylogenetic tree of Zot protein sequences from phage-WC36-1 and some related *Tubulavirales* phages. *Escherichia coli* was used as the outgroup. Bootstrap values (%) > 50 (A) and >80 (B) are indicated at the base of each node with the gray dots (expressed as percentages of 1000 replications). The accession numbers of phages encoding single-stranded DNA-binding proteins and Zot proteins are given after the phage names.

**Figure 5—figure supplement 3. Maximum likelihood phylogenetic tree of terL protein sequences from Phage-WC36-2, Phage-zth2-2, and some related *Caudoviricetes* phages.** *Cytomegalovirus* phage AD169 was used as the outgroup. Bootstrap values (%) > 80 are indicated at the base of each node with the gray dots (expressed as percentages of 1000 replications). The accession numbers of phages encoding terL proteins are given after the names of the phages.
