## Supplementary figures and images for "Polysaccharides induce deep-sea *Lentisphaerae* strains to release chronic bacteriophages"

### Figure 2 supplmental figure 1

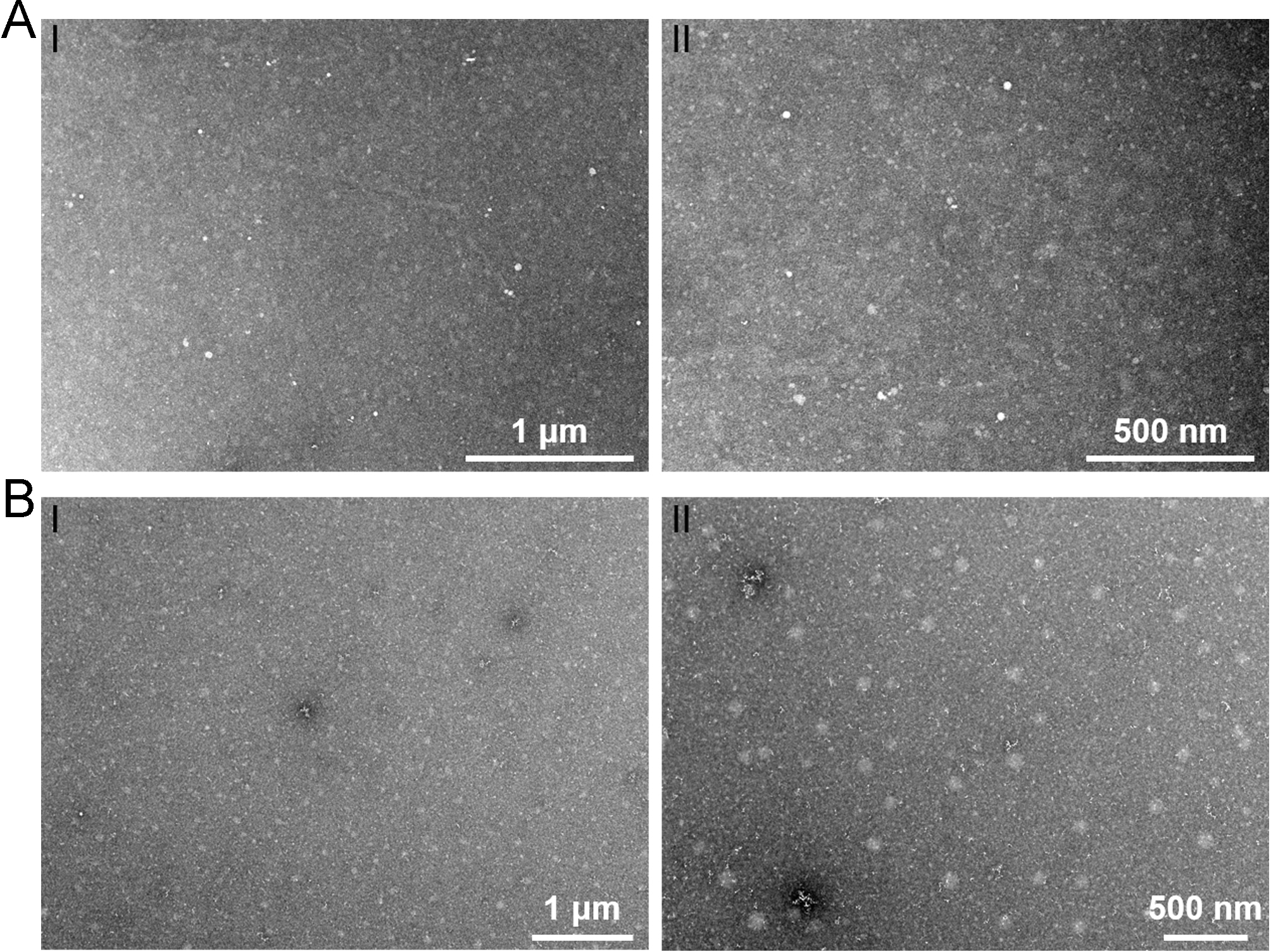

### Figure 2 supplmental figure 2

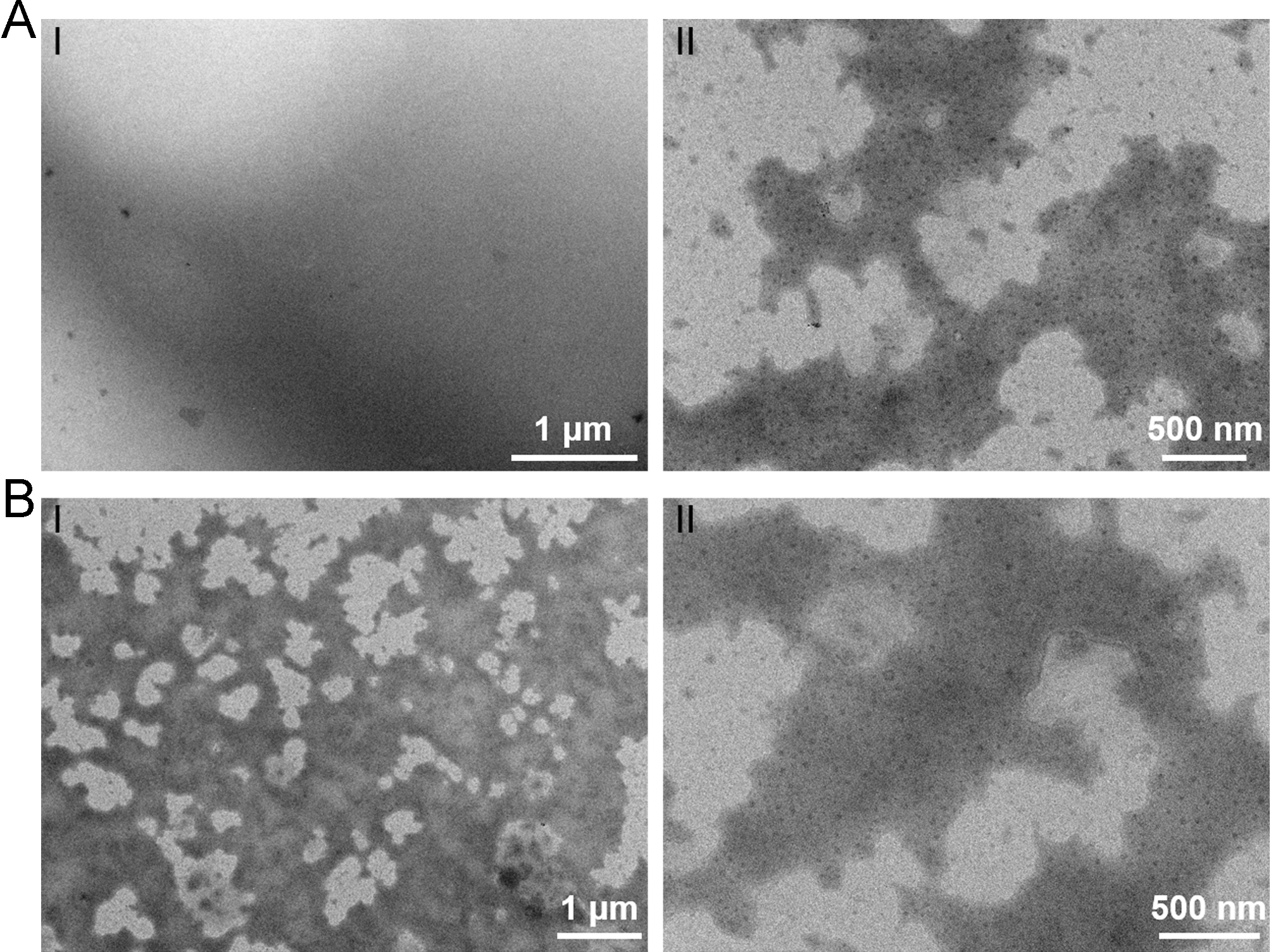

### Figure 2 supplmental figure 3

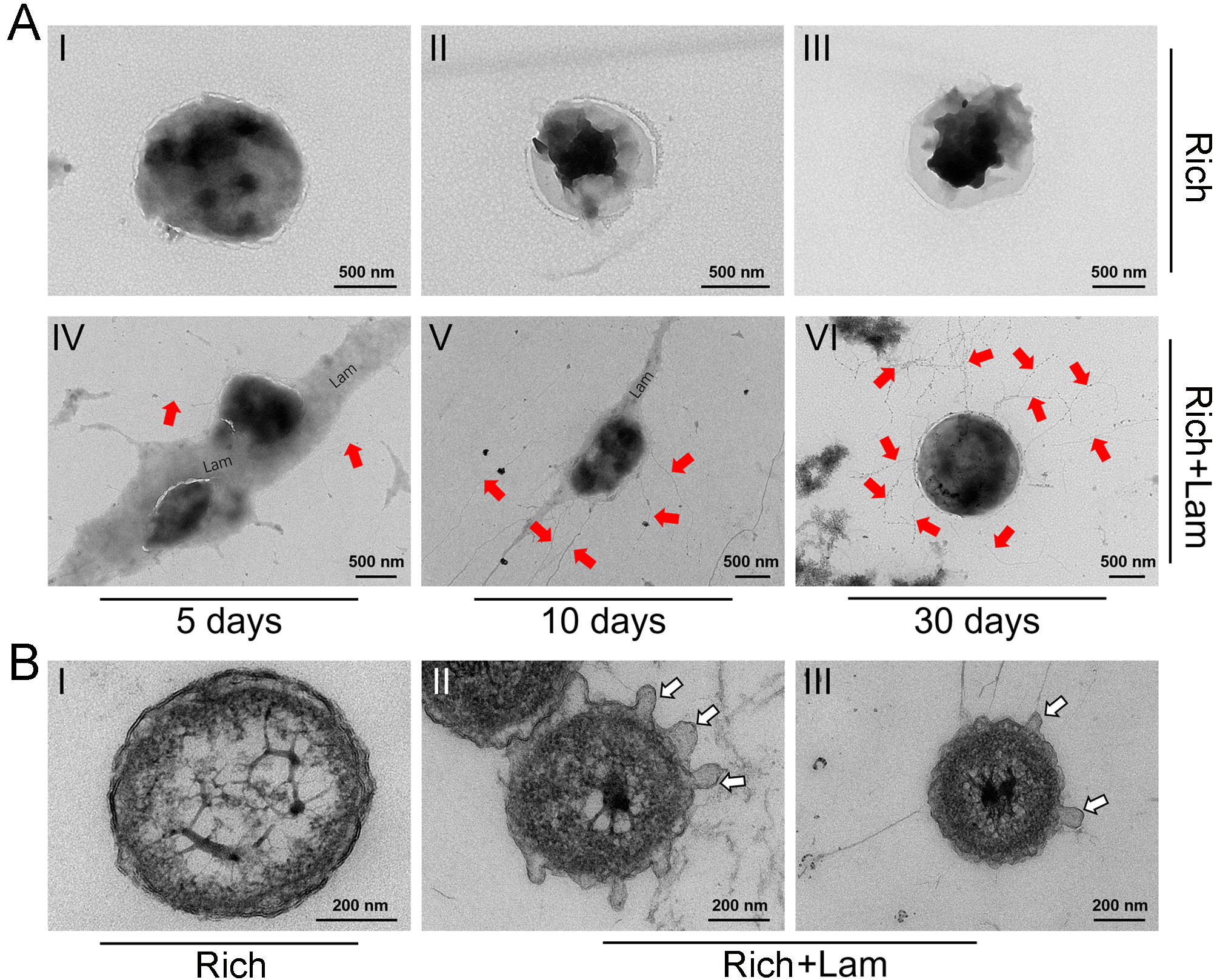

### Figure 3 supplmental figure 1

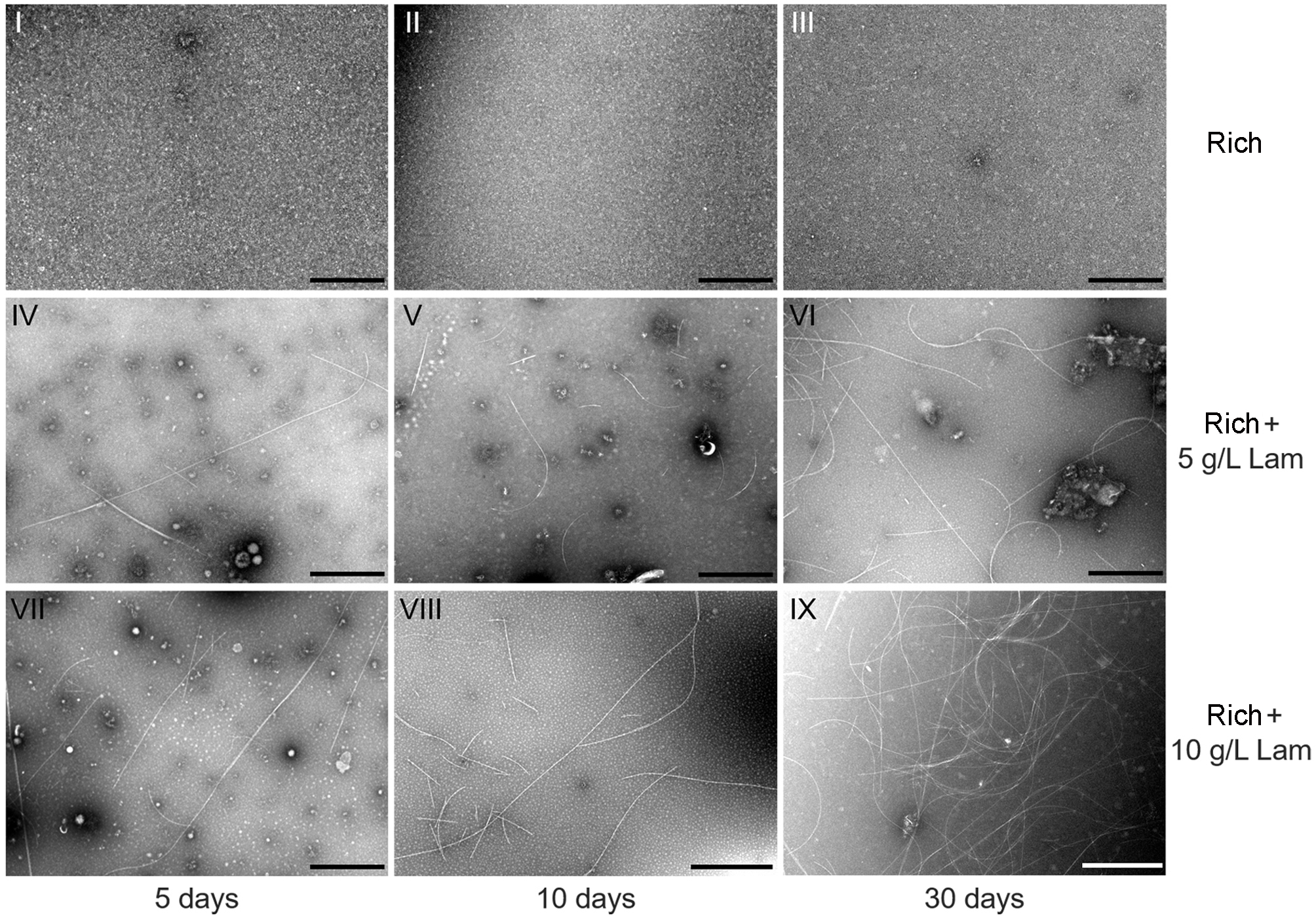

### Figure 4 supplmental figure 1

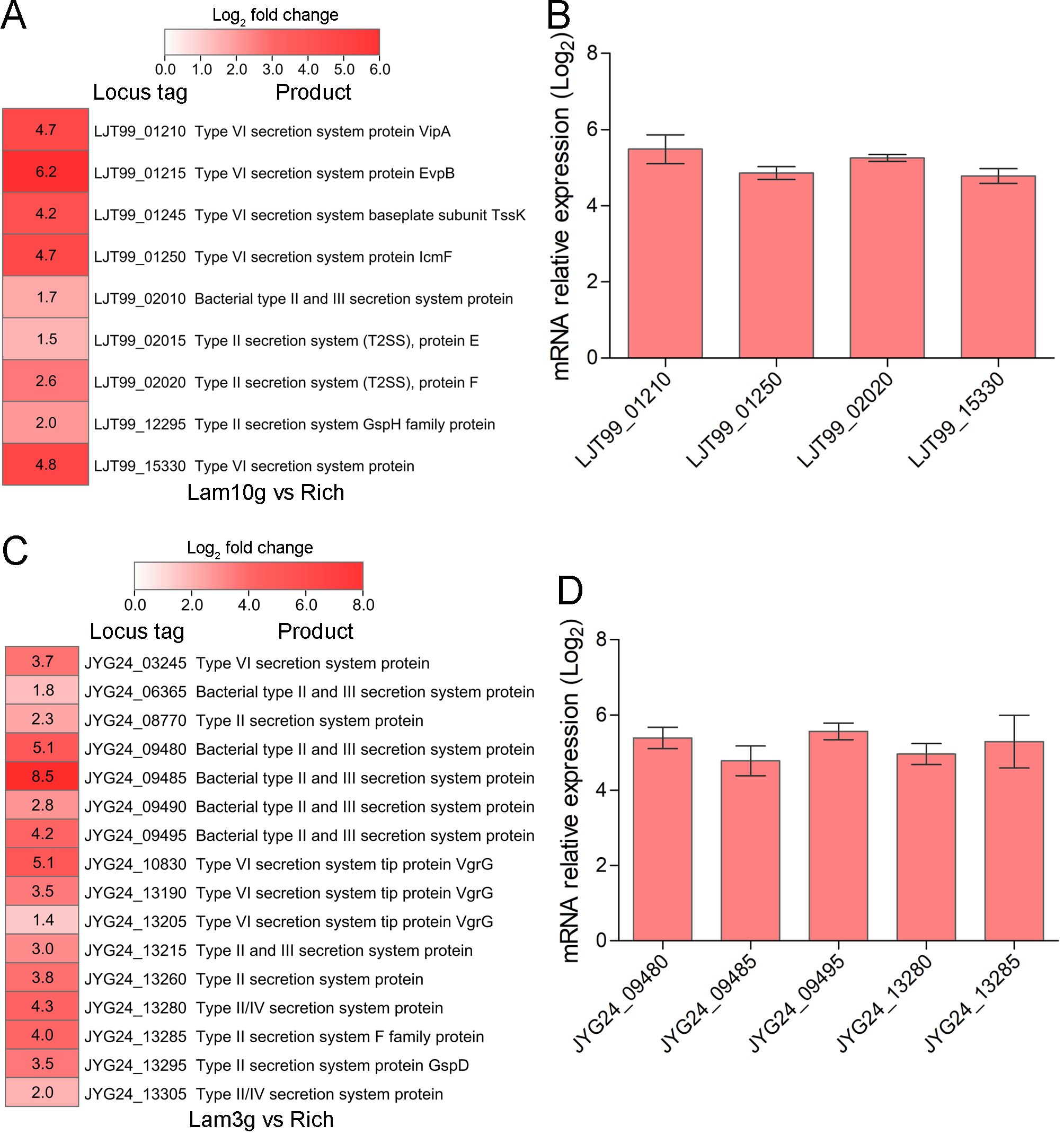

### Figure 5 supplmental figure 1

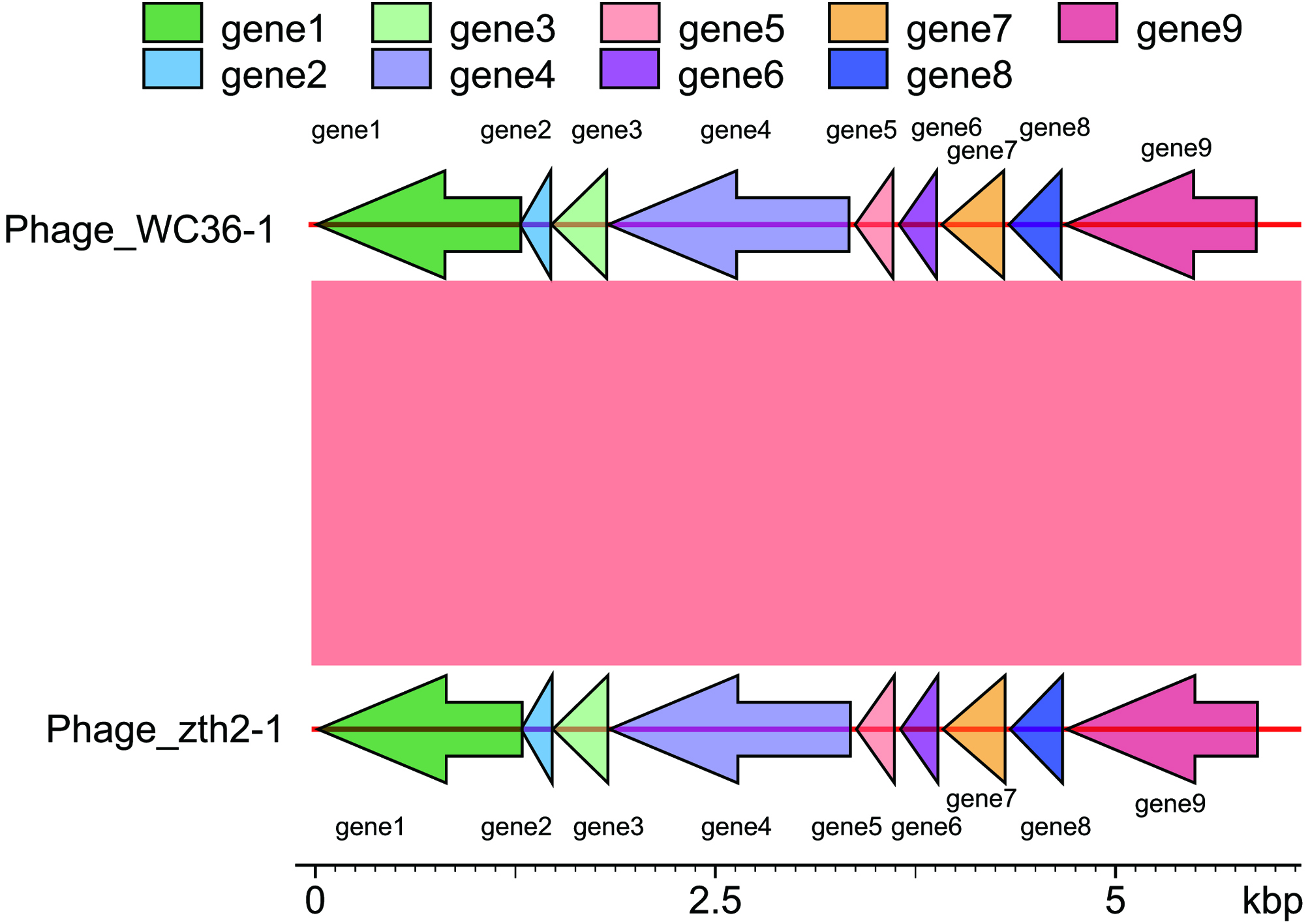

### Figure 5 supplmental figure 2

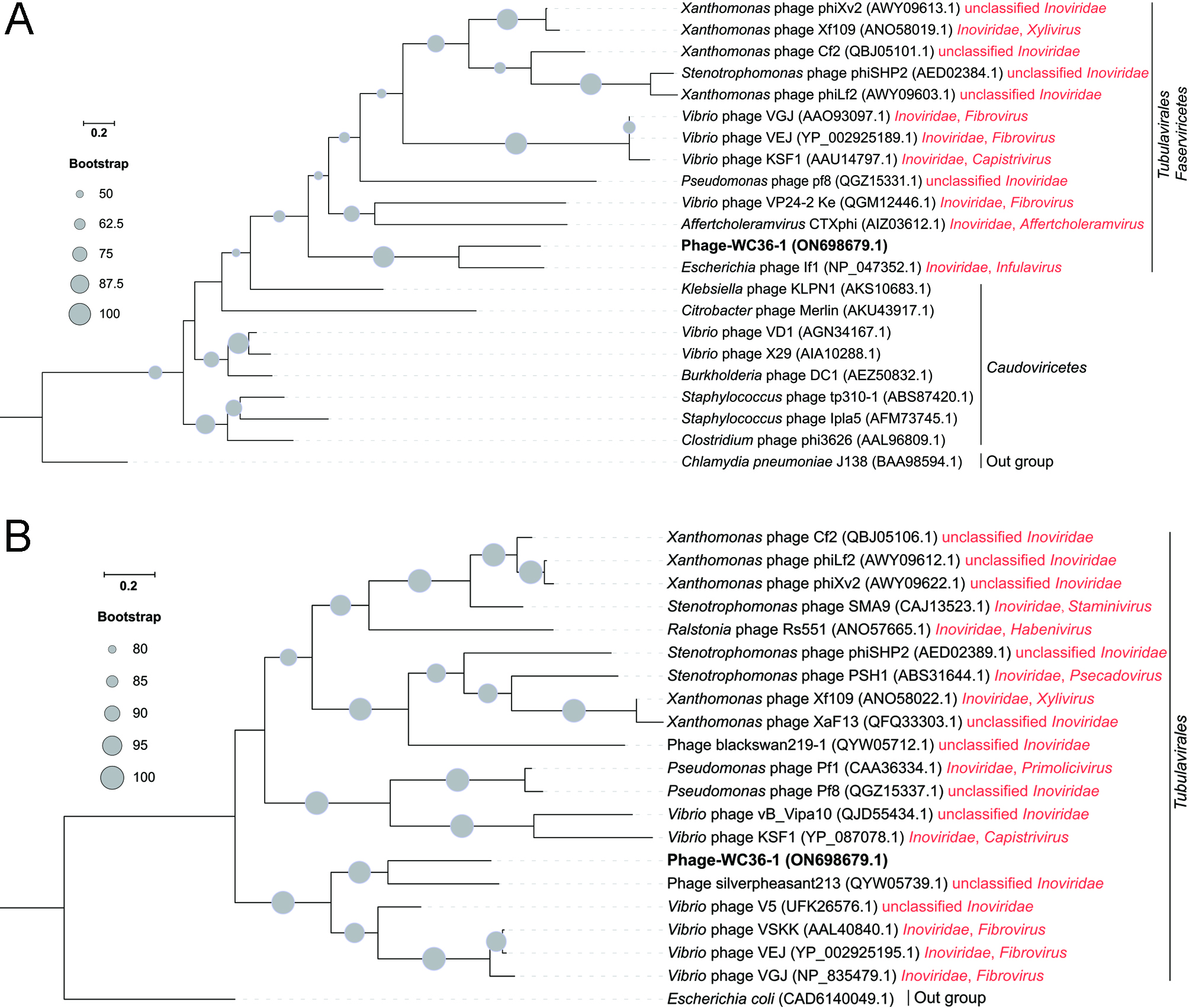

### Figure 5 supplmental figure 3

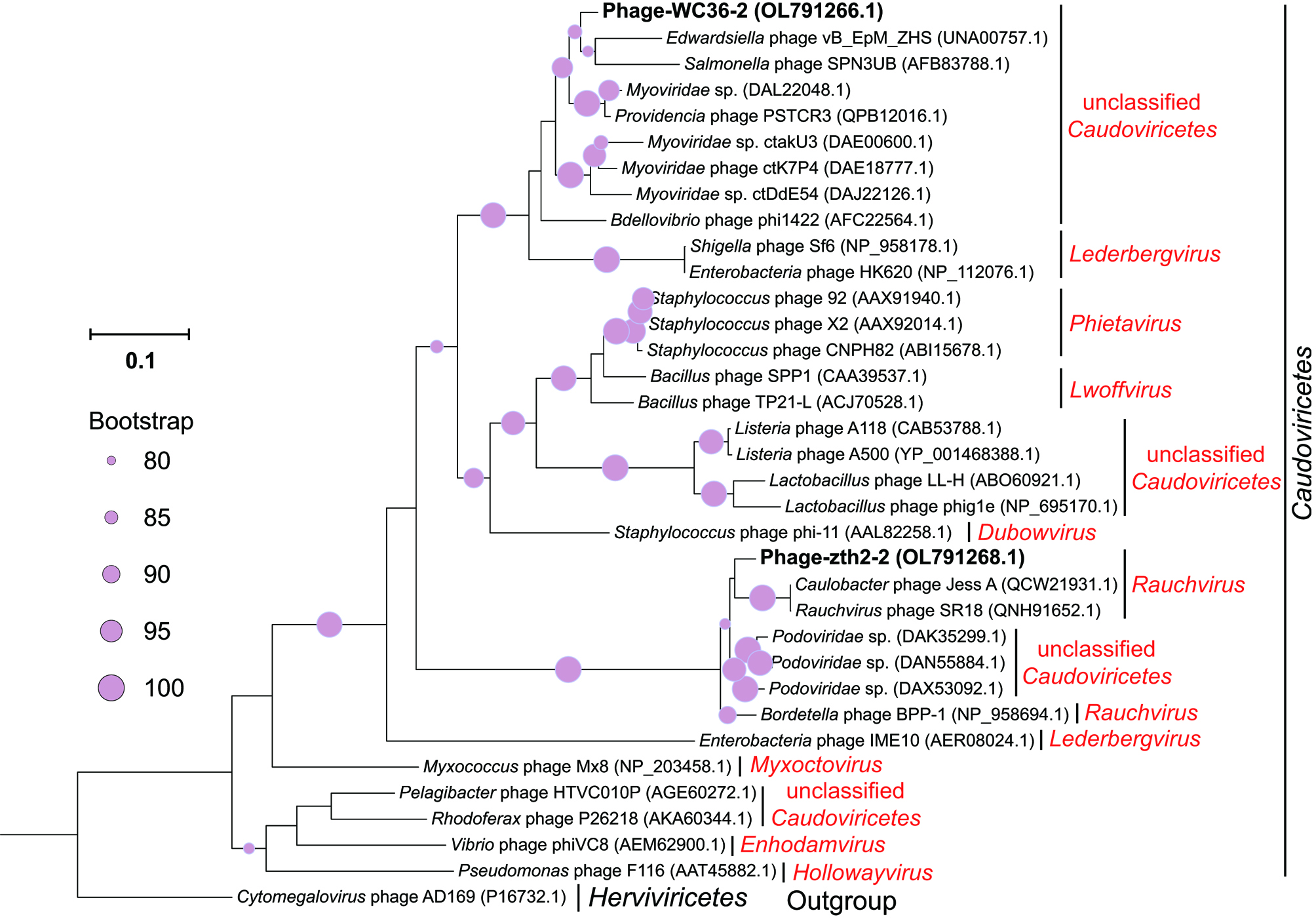
